# Pathway-specific asymmetries between ON and OFF visual signals

**DOI:** 10.1101/384891

**Authors:** Sneha Ravi, Daniel Ahn, Martin Greschner, E.J Chichilnisky, Greg D. Field

## Abstract

Visual processing is largely organized into ON and OFF pathways that signal stimulus increments and decrements, respectively. These pathways exhibit natural pairings based on morphological and physiological similarities, such as ON and OFF alpha ganglion cells in the mammalian retina. Several studies have noted asymmetries in the properties of ON and OFF pathways. For example, the spatial receptive fields (RFs) of OFF alpha cells are systematically smaller than ON alpha cells. Analysis of natural scenes suggests these asymmetries are optimal for visual encoding. To test the generality of ON-OFF asymmetries, we measured the spatiotemporal RF properties of multiple RGC types in rat retina. Through a quantitative and serial classification, we identified three functional pairs of ON and OFF RGCs. We analyzed the structure of their RFs and compared spatial integration, temporal integration, and gain across ON and OFF pairs. Similar to previous results from cat and primate, RGC types with larger spatial RFs exhibited briefer temporal integration and higher gain. However, each pair of ON and OFF RGC types exhibited distinct asymmetric relationships between receptive field properties, some of which were opposite to previous reports. These results reveal the functional organization of six RGC types in the rodent retina and indicate that ON-OFF asymmetries are pathway specific.

**Significance Statement:** Circuits that process sensory input frequently process increments separately from decrements, so called ‘ON’ and ‘OFF’ responses. Theoretical studies indicate this separation, and associated asymmetries in ON and OFF pathways, may be beneficial for encoding natural stimuli. However, the generality of ON and OFF pathway asymmetries has not been tested. Here we compare the functional properties of three distinct pairs of ON and OFF pathways in the rodent retina and show their asymmetries are pathway specific. These results provide a new view on the partitioning of vision across diverse ON and OFF signaling pathways

## Introduction

The division of sensory signals across neurons that respond to stimulus increments (ON) or decrements (OFF) is a common processing motif. Examples abound: olfactory receptor neurons in the cockroach respond to either increments or decrements in odor concentration (Burgstaller and Tichy, 2011); neurons in auditory cortex respond to increments or decrements of sound intensity (Scholl et al., 2017); neurons in the fish electrosensory system signal increasing or decreasing contrasts in amplitude modulations of an electromagnetic field (Berman and Maler, 1998; Clarke et al., 2014); and neurons from retina to visual cortex respond to increments or decrements of light intensity (Hartline, 1938; Hubel and Wiesel, 1962). Thus, understanding how and why ON and OFF pathways partition sensory input is central to an understanding of sensory processing.

In vision, the division of sensory processing between ON and OFF pathways is elaborate. The division originates at the first retinal synapse between photoreceptors and bipolar cells. Within one additional synaptic layer, the retina partitions visual scenes into 30-40 different channels, each instantiated by a distinct retinal ganglion cell (RGC) type (Field and Chichilnisky, 2007; Sanes and Masland, 2015). Many of these RGC types respond to either increments or decrements of light in their receptive field (RF) center (Hartline, 1938; Kuffler, 1953; Wassle and Boycott, 1991). Furthermore, many of these ON and OFF RGC types form pairs, such as ON and OFF alpha cells in cats and other mammals (Cleland and Levick, 1974; Cleland et al., 1975; Watanabe and Rodieck, 1989; Wassle and Boycott, 1991). These pairings have been established on both morphological and functional grounds. Morphologically, these pairs have dendritic fields that are similar in size and branching patterns, but that ramify in different depths of the inner plexiform layer (Wassle and Boycott, 1991; Dacey, 2004). Functionally, these pairs exhibit similar receptive fields with a polarity reversal. However, multi-neuron measurements have identified systematic ‘asymmetries’ between some paired ON and OFF RGC types (Chichilnisky and Kalmar, 2002; Ratliff et al., 2010). For example, both ON parasol RGCs exhibit larger spatial RFs than their OFF-cell counterparts. Asymmetries between ON and OFF pathways have also been observed in temporal integration, contrast response functions, absolute sensitivity, nonlinear spatial integration, and adaptation (Chichilnisky and Kalmar, 2002; Nirenberg et al., 2010; Pandarinath et al., 2010; Ala-Laurila and Rieke, 2014; Turner and Rieke, 2016).

These asymmetries have been studied mostly in alpha and parasol RGCs, which are probably homologs (Crook et al., 2008a). This raises the question, how ubiquitous are these asymmetries? Analysis of natural scenes suggests that RF size asymmetries may be an efficient coding scheme for natural scenes (Ratliff et al., 2010)(Barlow, 1961; Pandarinath et al., 2010; Karklin and Simoncelli, 2011). These results suggest asymmetries may be preserved across ON and OFF pathway pairs. However, these analyses were agnostic to the particular aspects of the visual image represented by distinct cell types, which may dictate distinct asymmetries (or even symmetry) for efficient coding.

The goal of this study was to measure the organization of RFs across multiple pairs of ON and OFF RGCs to determine the extent to which asymmetries are general or pathway specific. We measured the RF properties of hundreds of simultaneously recorded rat RGCs using a multi-electrode array. We developed a procedure for functionally classifying RGCs based on their responses to diverse visual stimuli. This classification yielded six irreducible cell types -- three pairs of ON and OFF RGC types. Across three pairs of ON and OFF RGCs from these six types, we found that the relative organization and the presence of functional asymmetries was pathway dependent. Each pair exhibited a distinct set of asymmetries in spatiotemporal integration and contrast response functions. These results indicate that asymmetries between ON and OFF pairs are common, but that the differences between pairs vary with the cell type and their light response properties.

## Materials and Methods

### Tissue preparation and MEA Recordings

All experiments followed procedures approved by the Institutional Animal Care and Use Committee of Duke University and Salk Institute for Biological Studies. Long Evans rats were euthanized by IP injection of ketamine and xylazine. Retinas were removed in darkness under infrared illumination with infrared converters as described previously (Anishchenko et al., 2010; Yu et al., 2017). A ~1.5 x 3 mm segment of dorsal retina centered 3.5-4 mm above the optic nerve and +/- 1mm along the vertical meridian was isolated. This region of retina was targeted to minimize variability across experiments and to target retinal locations with cones expressing mostly M-opsin. The retina was placed RGC side down on an electrode arrays consisting of 512 electrodes at 60 µm interelectrode spacing, spanning an area of 0.9 x 1.8 mm (Litke et al., 2004). The voltage trace recorded on each electrode was bandpass filtered between 80 and 2,000 Hz, sampled at 20 kHz, and stored for off-line analysis (Frechette et al., 2005). Spikes were initially sorted by an automated algorithm and the resulting clusters were checked and corrected manually using custom spike sorting software (Shlens et al., 2006; Yu et al., 2017). The autocorrelation function of sorted spikes was used to validate putative RGCs by checking for a refractory period (1.5 ms (Field et al., 2007)). To track the RGCs across different visual stimuli, spike shapes were sorted in the same subspace determined by principal components analysis (PCA) of the spike waveforms. Neuron identity was further confirmed across different stimuli by checking that the electrical image (EI (Petrusca et al., 2007)) for each neuron matched across conditions. A matched neuron between two stimulus conditions was determined by the EI pair with the highest inner product across the two stimulus conditions (Field et al., 2009). A typical experiment resulted in recording and tracking the responses of 300-400 RGCs across three visual stimuli.

### Visual Stimuli and RGC Response Properties

Visual stimuli from a gamma-corrected CRT video display (Sony Trinitron) refreshing at 120 Hz, or an OLED display (Emagine) refreshing at 60 Hz, were focused on the retina via an inverted microscope (Yu et al., 2017). Two different stimuli were used to measure the functional properties of recorded RGCs; each was photopic with a mean intensity of either between 3000 or 10,000 photoisomerizations/rod/s (Field et al., 2009; Yu et al., 2017). First, a checkerboard noise stimulus was used to estimate the spatiotemporal RF by reverse correlation (Chichilnisky, 2001). Each checker of the noise stimulus was 40×40 microns on the retina and noise images were updated at 60 Hz. Second, sine wave gratings with a spatial period of 320 µm on the retina were drifted in 8 directions at two speeds (150 and 600 µm/s). This stimulus identified RGCs that were sensitive to motion (Figure 1A) (Yu et al., 2017).

**Figure 1:**
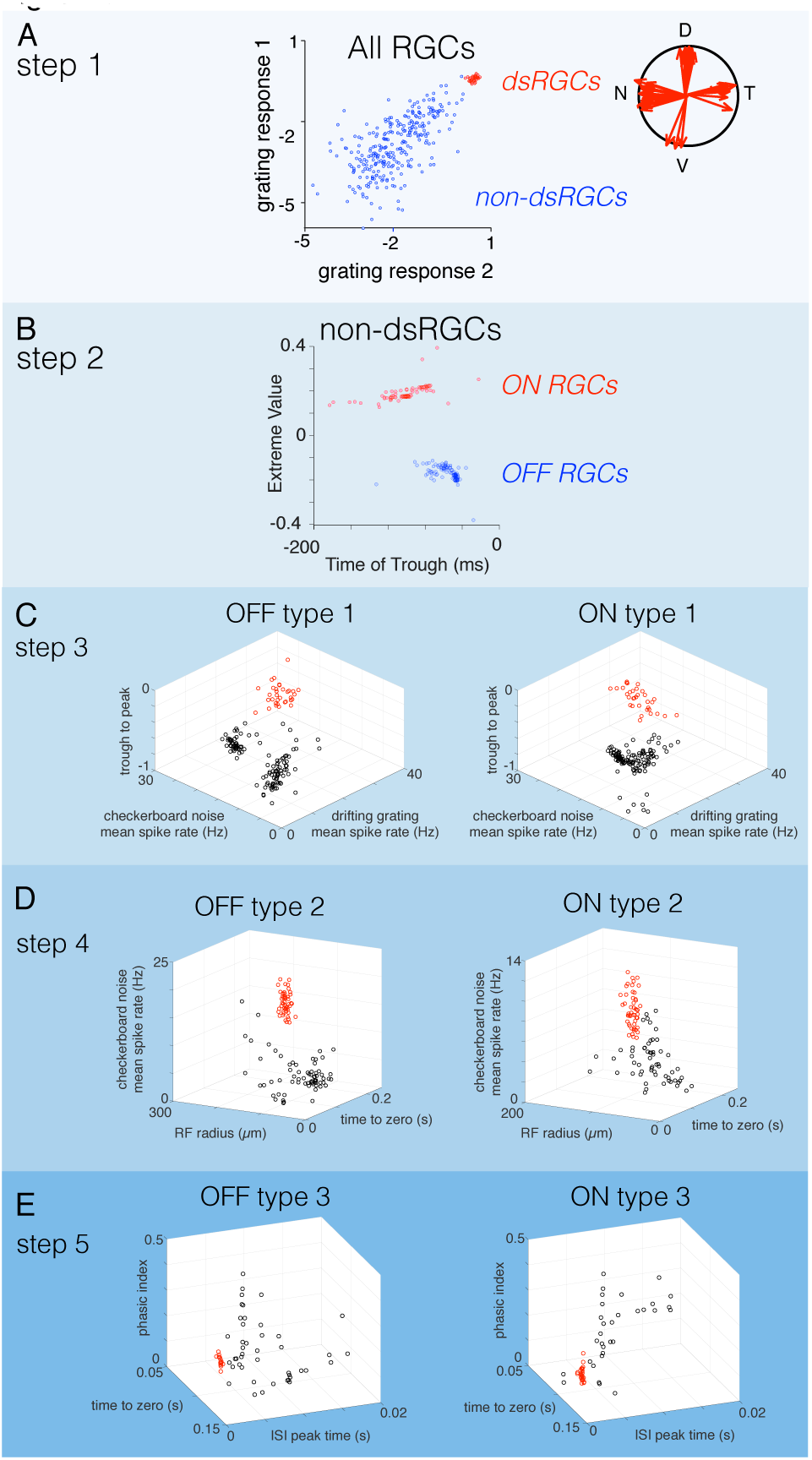
Serial Classification of RGCs yield three pairs of ON and OFF cells. **A.** In step 1 of the classification, direction selective RGCs are segregated from all other cells based on their responses to drifting gratings. Grating response 1 and 2 are the natural log of the vector magnitude to a grating with a spatial period of 320 µm drifting at 150 and 600 µm/s, respectively. Gratings were drifted in 8 directions to estimate the vector magnitudes of their tuning. **B.** In step 2, ON and OFF RGCs were segregated by the value of the extrema and time to trough of their temporal RFs estimated from their STA. **C.** In step 3, a pair of ON and OFF RGCs (red points) were classified from all other ON and OFF cells, respectively. The parameter spaces used to classify these two types were identical and consisted of the mean spike rates to checkerboard noise (stixel size 40×40 µm, 60 Hz refresh) and a drifting grating (spatial period 320 µm, speed, 150 µm/s), as well as the trough to peak ratio of their temporal RFs. **D.** In step 4, ON and OFF RGCs identified in step 3 were removed, and the remaining ON and OFF RGCs were classified in a new parameter space defined by the mean spike rate to checkerboard noise, RF radius, and the time to zero of the temporal RF. **E.** In step 5, ON and OFF RGCs identified in the two previous steps were removed and the remaining ON and OFF cells were classified in a new parameter space defined by the phasic index (estimated from the temporal RF, see Materials and Methods), time to zero of the temporal RF, and the peak time of the interspike-interval (ISI) distribution. At each step of the classification, groups of cells were distinguished by a two-Gaussian mixture model.

### RGC classification

RGCs from seven retinas were classified in this study. The number of cells identified for each type in each retina are provided in Table 1. The classification approach consisted of two stages: a feature selection process followed by a serial, quantitative classification using unsupervised learning. The feature selection process identified response properties that robustly isolated one or a small number of RGC types from all other types (e.g. isolating DS-RGCs from nonDS-RGCs, Figure 1A). The quantitative classification clustered neurons using these features by a two-Gaussian mixture model.

**Table 1.** RGC counts are provided for the six RGC types identified and examined across the seven retinal recordings used in this study.

| **Retina:** | **1** | **2** | **3** | **4** | **5** | **6** | **7** |
| --- | --- | --- | --- | --- | --- | --- | --- |
| *ON brisk sustained* | 33 | 24 | 20 | 26 | 22 | 25 | 38 |
| *ON brisk transient* | 51 | 40 | 50 | 39 | 48 | 53 | 37 |
| *ON small transient* | 28 | 20 | 23 | 20 | 21 | 30 | 9 |
| *OFF brisk sustained* | 34 | 27 | 26 | 31 | 33 | 33 | 23 |
| *OFF brisk transient* | 52 | 30 | 44 | 36 | 37 | 40 | 48 |
| *OFF small transient* | 15 | 10 | 7 | 10 | 14 | 21 | 12 |

### Stage One

The feature selection process was performed using one of the seven retina recordings in this manuscript. This stage was used to identify response parameters that distinguished one set of RGCs from all others. In this initial dataset, high-dimensional data was parameterized and visualized in a lower dimensional space by PCA. These spaces consisted of either two or three dimensions, each defined by a response parameter such as the overall spike rate or the shape of the temporal RF (e.g. Figure 1C). Limiting the dimensionality facilitated robustly clustering RGCs with relatively limited data (e.g. a few hundred RGCs). Once a set of response features were identified that clearly separated one group of RGCs from the others, the spatial RFs of the grouped RGC were inspected to check whether they were regularly spaced. If grouped RGCs were regularly spaced, the features used were saved for quantitative clustering (see Stage Two). Performing feature selection before quantitative classification improved the performance of the unsupervised clustering algorithm by minimizing misclassification rates.

### Stage Two

To quantitatively cluster each group of RGCs (Figure 1), a two Gaussian mixture model (GMM) was fit in the same two or three-dimensional feature space defined above in Stage One. The GMM allowed boundaries to be drawn between clusters according to the maximum likelihood that RGCs belonged to one Gaussian distribution or the other. RGC types were classified one at a time in a serial fashion to prevent overfitting and avoid ambiguity in choosing the right number of clusters. Each cluster was tested for statistical significance (Tukey’s range test), and the irreducibility of each type was verified by testing for a mosaic organization (Figure 3). The order of this serial classification and the response parameters that consistently identified RGCs across recordings is shown in Figure 1.

**Figure 2.**
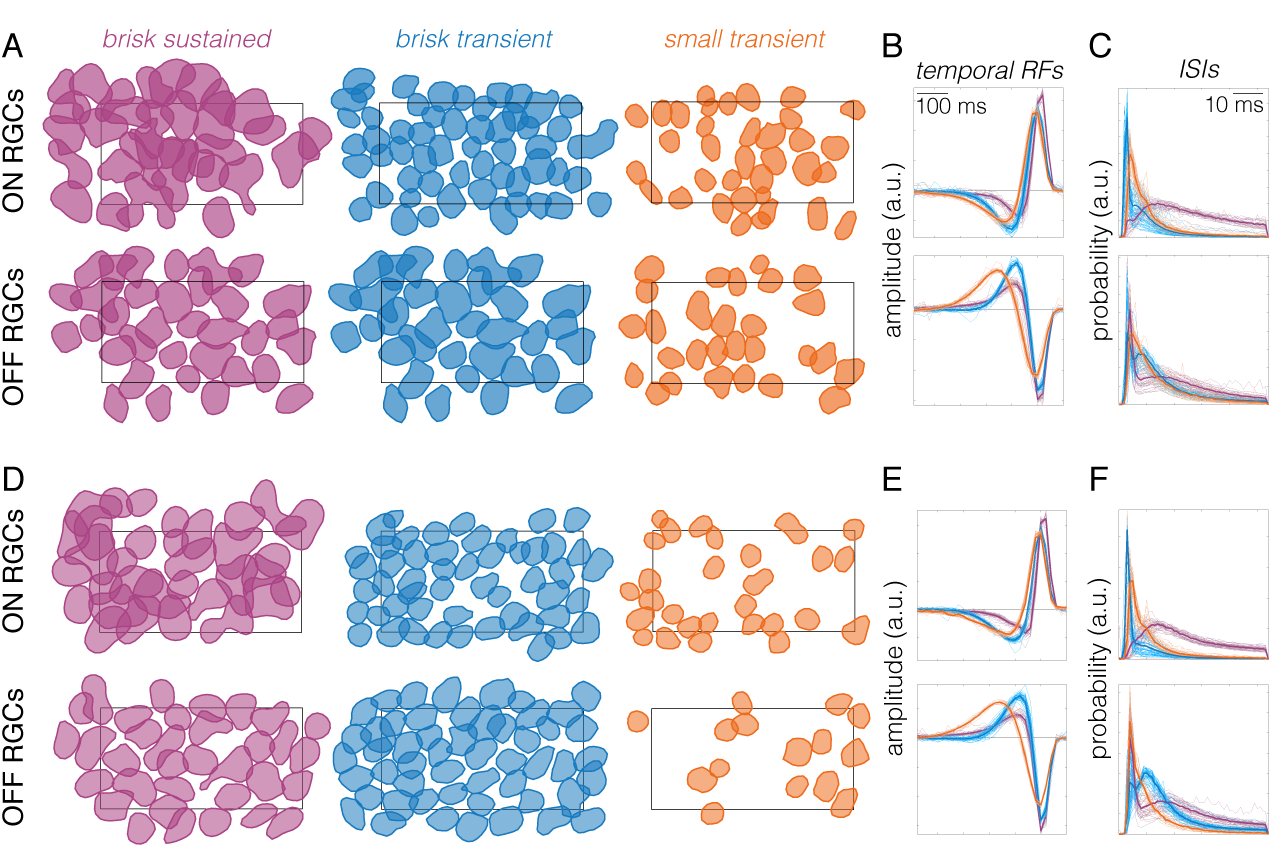
Classified ON and OFF RGCs exhibit a mosaic-like organization. **A.** Spatial RFs of ON and OFF brisk sustained (purple), brisk transient (blue) and small transient (orange) RGCs identified in one retina. Spatial RFs are shown as a contour plotted at 0.6065 of the peak amplitude (equivalent to 1 SD of Gaussian). Rectangle shows the outline of the MEA (900 x 1800 µm). **B.** Temporal RFs of all cells shown in **A**, with ON cells on top and OFF cells on bottom. Thin lines are individual cells, thick lines are mean. Color conventions same as A. **C.** Inter-spike interval (ISI) distributions for all cells in **A**. Color and line conventions same as **A** and **B**. **D-F.** Same as **A-C**, but for a second retina.

**Figure 3.**
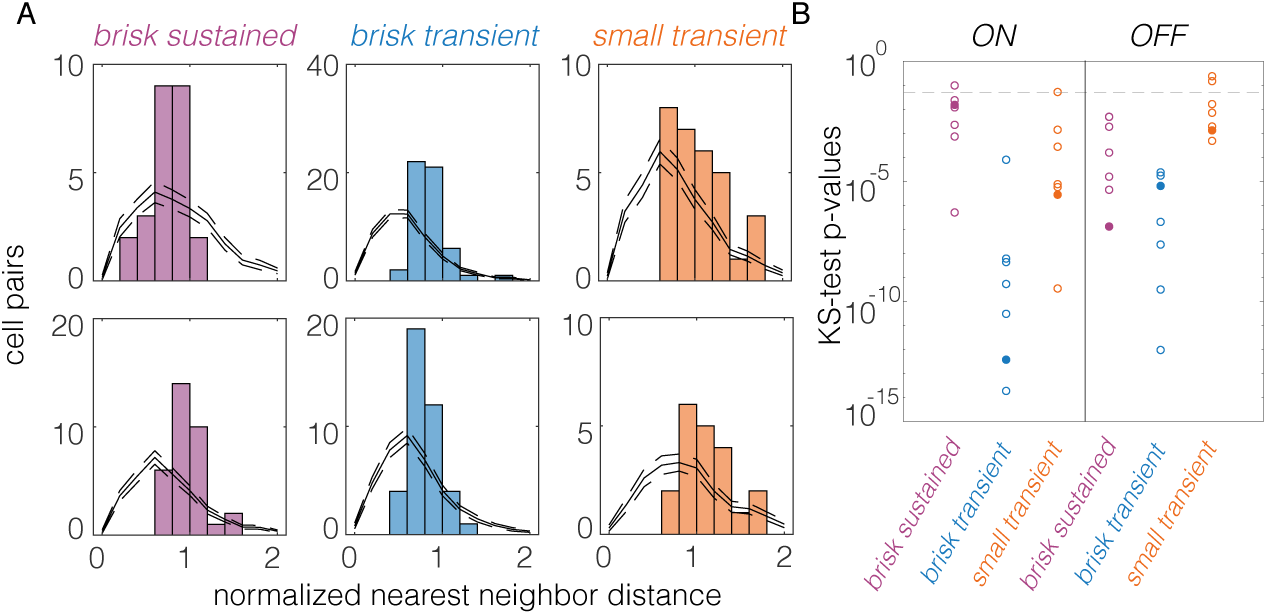
Normalized nearest neighbor distributions (NNNDs) indicate mosaic-like arrangement of spatial RFs. **A.** NNNDs for brisk sustained, brisk transient, and small transient cells, ON cells are top, OFF cells are bottom. Data are from one retina. Black lines show expected NNNDs for randomly sampled cell locations (see Materials and Methods); dashed lines show 95% CI. **B.** P-values from a two-sample Kolmogorov-Smirnov (KS) test for observed NNNDs arising from random cell locations. Fill circles correspond to data shown in **A**.

### Verifying RGC Types

Clustered RGCs were identified as an irreducible cell type by inspecting the normalized nearest neighbor distribution (NNND; Figure 3) (DeVries and Baylor, 1995; Field et al., 2007). The NNND is defined as *2R / (S1 + S2)*. *R* is the distance between the spatial RF of each RGC and its nearest neighbor’s RF. *S1* and *S2* are SDs of the Gaussian fits for each RGC’s spatial RF measured along the line connecting the centroids. If the two spatial RF ‘touch’ at the 1-SD contour for each cell, then the NNND will equal 2.

NNNDs indicate a mosaic-like arrangement of RFs when they exhibit a clear exclusion zone at short nearest-neighbors at distances (Wassle and Riemann, 1978). To test the null hypothesis that the observed NNND were consistent with a random sampling of RGCs, we generated 100 NNND distributions from randomly sampled RGCs within each experiment (Figure 3A). The number of sampled cells equaled the number of RGCs in the original mosaic. A two-sample Kolmogorov-Smirnov test was used to estimate the probability that the observed NNND was consistent with that expected from a randomly sampled set of RGCs. In 38 of 42 mosaics tested, the null hypothesis was rejected with p < 0.05 (Figure 3B).

### Estimation of linear spatiotemporal RFs

A linear approximation to the spatiotemporal RF of each RGC was obtained by reverse correlation to compute the STA (Chichilnisky, 2001). Frames up to 500 ms preceding a spike were included in the analysis. The spatial RF was the set of stimulus pixels (stixels) whose absolute peak intensity exceeded 4.5 robust standard deviations of all pixel intensities (Yu et al., 2017). The temporal RF was defined as the time-dependent average of these significant stimulus pixels. Once the temporal RF was computed, the dot product between every stixel of the STA was computed with the temporal RF. This collapsed the STA across time to a single image, which was used as an estimate of the spatial RF.

This analysis to extract estimates of the spatial and temporal RFs assumes the spatiotemporal RF is separable into a single spatial and temporal filter. The validity of this assumption was examined using singular value decomposition (SVD; (Golomb et al., 1994)). SVD factorizes a matrix into a rank-ordered set of vector pairs whose outer products are weighted and linearly combined to reproduce the original matrix. A perfectly space-time separable RF will produce a single pair of non-zero vectors capturing the spatial and temporal RFs respectively. Prior to performing SVD, a Gaussian spatial filter was applied to the full spatiotemporal RF to reduce noise in the STA. This Gaussian filter was circular with an SD of 0.75 stixels. After applying this filter, SVD indicated that across cell types, >90% of the variance in the STA could be captured by the outer product of a single pair of spatial and temporal filters. This indicates that the linear RF structure was largely consistent with a space-time separable model.

### Space-time plots

To generate average space-time plots of RGC RFs (Figure 5), the entire spatiotemporal RF was filtered for each cell with a circular Gaussian filter, SD = 0.75 stixels. A 21×21 (924 microns x 924 microns) stixel region around the center of mass of the spatial RF was cropped. The average 3-dimensional spatiotemporal RF of each RGC type was computed by averaging together all the cropped and filtered spatiotemporal RFs of all cells of that type across all recordings. The 3-dimensional spatiotemporal RF was collapsed to 2 dimensions by extracting the intensities along one spatial axis.

**Figure 4.**
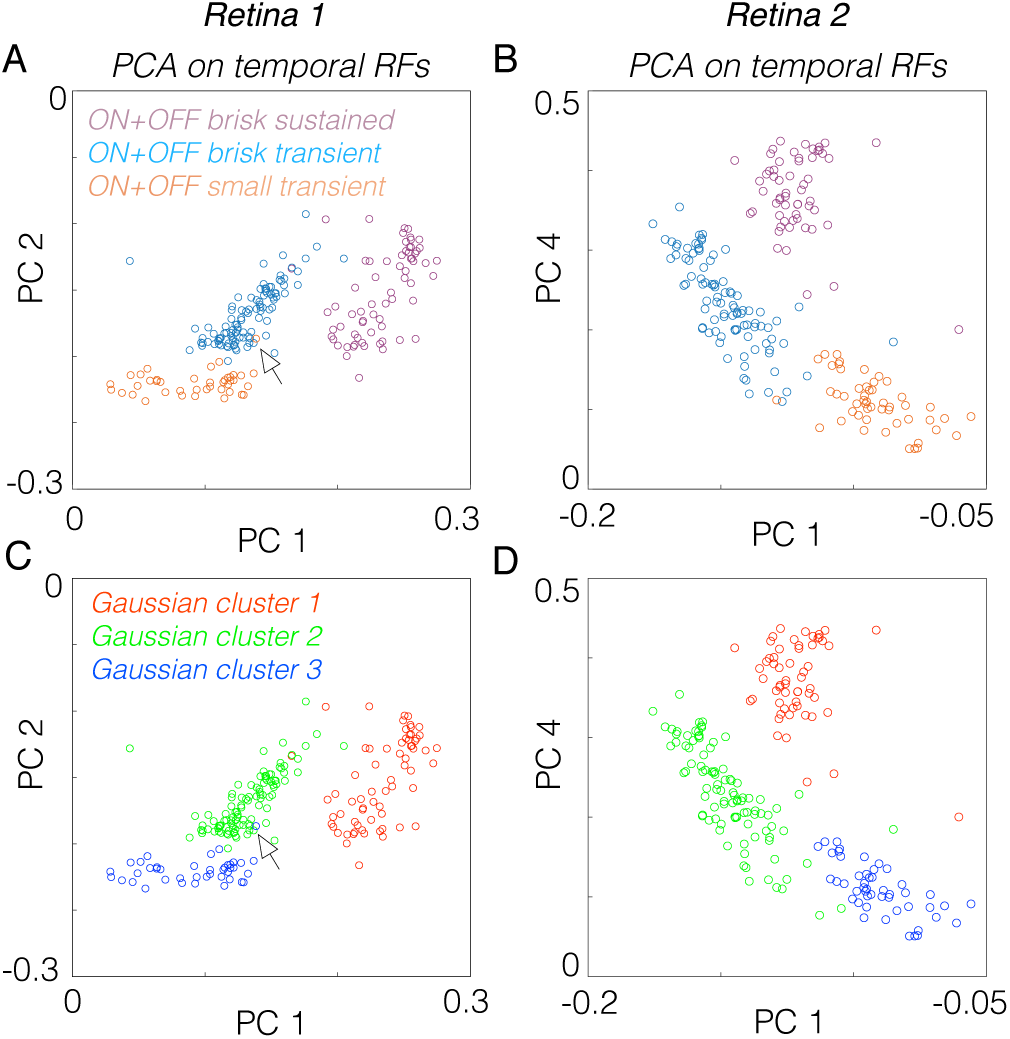
Temporal RFs of ON and OFF pairs cluster together after accounting for polarity differences. **A.** PCA applied to the temporal RFs of brisk sustained, brisk transient, and small transient cells from one experiment. The temporal RFs of OFF cells were multiplied by -1 to invert their polarity prior to PCA. Each circle represents one RGC, circles were colored by cell type determined by the classification in Figure 1. **B.** Same analysis as A, but for a second retina and weights associated with PC 4 are plotted instead of PC 2. **C.** Same data as in A, but a three Gaussian mixture model was fit to the data in the space defined by the first 5 principal components, which captured >99% of the variance in the data. This fit finds the best 3-group description of the data (provided each group is well described by a multivariate Gaussian distribution). The Gaussian mixture model clustered the temporal RFs identically to the groupings defined by combining ON and OFF pair together. Even points that appear outside of their appropriate group (see arrowheads) in the two-dimensional plot are correctly classified by the Gaussian mixture

**Figure 5.**
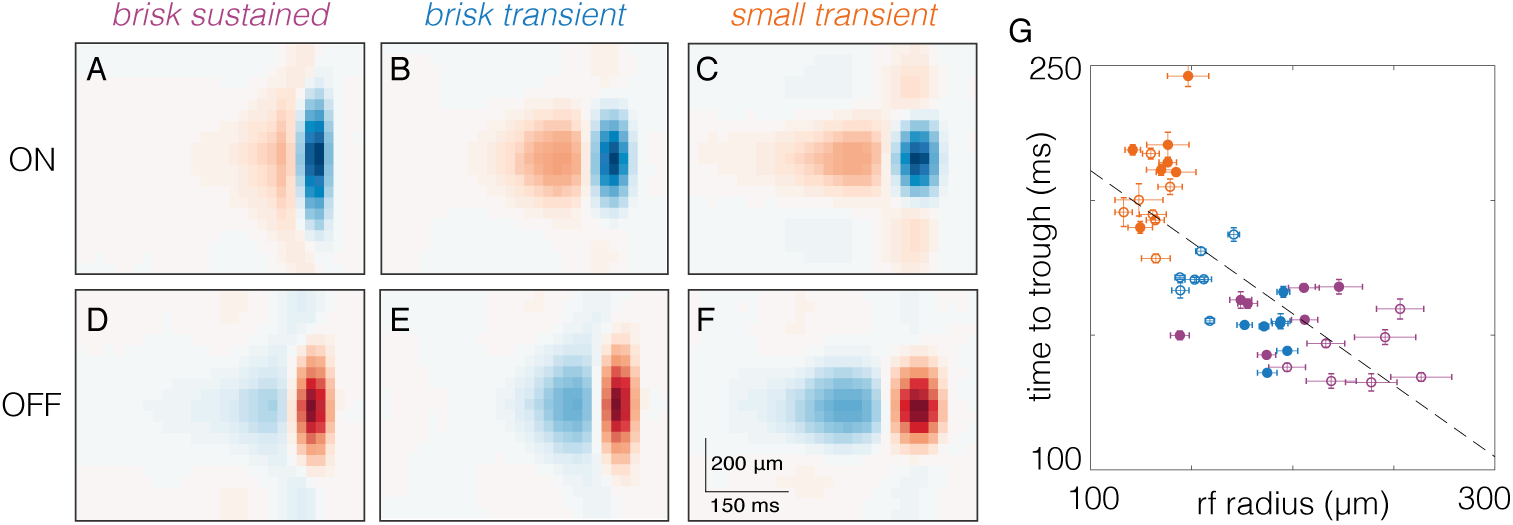
RGC types with larger spatial RFs exhibit briefer temporal RFs. **A-F.** Average spacetime RFs from one retina of ON (**A-C**) and OFF(**D-F**) brisk sustained, brisk transient, and small transient RGCs. **G.** Comparison of spatial integration (rf radius) to temporal integration (time to trough). Each point corresponds to one RGC type from one retina, filled (open) symbols are OFF (ON) RGCs. Brisk sustained are purple, brisk transient are blue, and small transient are orange. Dashed line is the best fit line to the data (slope = -0.531, y-intercept = 264.15 ms). Error bars are SE.

### Estimation of Contrast response functions

Contrast response functions were estimated from the static nonlinearity computed by convolving the spatiotemporal RFs with the checkerboard noise stimulus (Chichilnisky, 2001). This yielded an instantaneous generator signal for each frame of the stimulus that was used to generate a histogram of observed spike counts for each generator signal. This histogram was fit with a logistic function. The slope (b) and offset (a) were parameters from the logistic function fit to the SNL: (c/(1+exp(-b(x-a))). To check that the static nonlinearity was accurately fit, simulated spikes were generated from a model Linear-Nonlinear Poisson neuron in response to a checkerboard white noise stimulus. A logistic function was used in the simulation for the nonlinearity. When total spike counts were matched between simulated and real neurons, the model fitting produced estimates of the slope and offset within 1% of the values set in simulation.

### Accuracy of the LNP model

An important caveat in the RF measurements presented here is that they are linear estimates. These estimates have been shown in some circumstances to accurately capture the stimulus features that drive RGC spiking (Chichilnisky, 2001; Keat et al., 2001; Pillow et al., 2005). However, for some RGC types, stimulus features interact nonlinearly in space and/or time (Hochstein and Shapley, 1976; Schwartz et al., 2012; Freeman et al., 2015). To determine the capacity of these linear RF estimates and contrast response functions to capture the relationship between the stimulus and spiking, we cross-validated the model to a repeated checkerboard noise stimulus in a subset of experiments (retinas 2 and 3, Table 1). A 10 s checkerboard noise sequence (40×40 µm stixels, 60 Hz refresh) was repeated 100 times. For a given RGC, the LNP model generated from the spatiotemporal RF and static nonlinearities estimated from the non-repeating checkerboard noise was used to predict the response to the repeated checkerboard stimulus (not used in the original estimate of the STA or static nonlinearity). Across cells of all six types, spike trains generated by the LNP model captured 51-73% of the explainable variance (data not shown).

### Parameterizing stimulus responses

#### Vector sum for drifting gratings

The total spike count from RGCs to 8 presentations of a grating drifting in each of 8 directions was calculated and normalized by the maximum count. This yielded 8 vectors that had magnitudes ranging between 0 and 1. The sum of these vectors identified the preferred direction of the RGC (Elstrott et al., 2008; Rivlin-Etzion et al., 2012) and the magnitude of this vector was used to estimate the strength of tuning and classify dsRGCs from non-dsRGCs in Figure 1A. The vector sum was not normalized to 1 to allow the vector magnitude to range from zero to infinity. This allowed the Gaussian mixture model to be fit to the log (base 2) of the vector sum: these distributions were approximately log-normal.

#### Firing rate for drifting gratings and checkerboard noise

The firing rates in response to drifting gratings were calculated by dividing the total spike count by the number of stimulus repeats (8), directions (8) and length of time that the grating was presented to the retina (8 or 10 s). For checkerboard noise, the total number of spikes during the presentation of the checkerboard noise was divided by the total time.

#### Parameters of the temporal RF from checkerboard stimuli

The time-to-peak and time-to-trough were taken from the global maximum and minimum, respectively, in the temporal RF. The zero crossing was calculated as the time closest to the spike at which the temporal RF transitioned from positive to negative values for OFF cells and vice-versa for ON cells. The maximum and minimum values were taken as the global maximum and minimum in the temporal RF, respectively. A phasic index (PI) was calculated from the temporal RF as the absolute value of the sum of the positive and negative areas divided by the sum of their absolute values (e.g. |(a+b)| / (|a| +|b|)). The PI ranges from zero to one: zero corresponds to a biphasic temporal RF with the area above and below zero being equal; one corresponds to a monophasic temporal RF. The biphasic index (Figure 6D) equaled 1 – PI (Petrusca et al., 2007).

**Figure 6.**
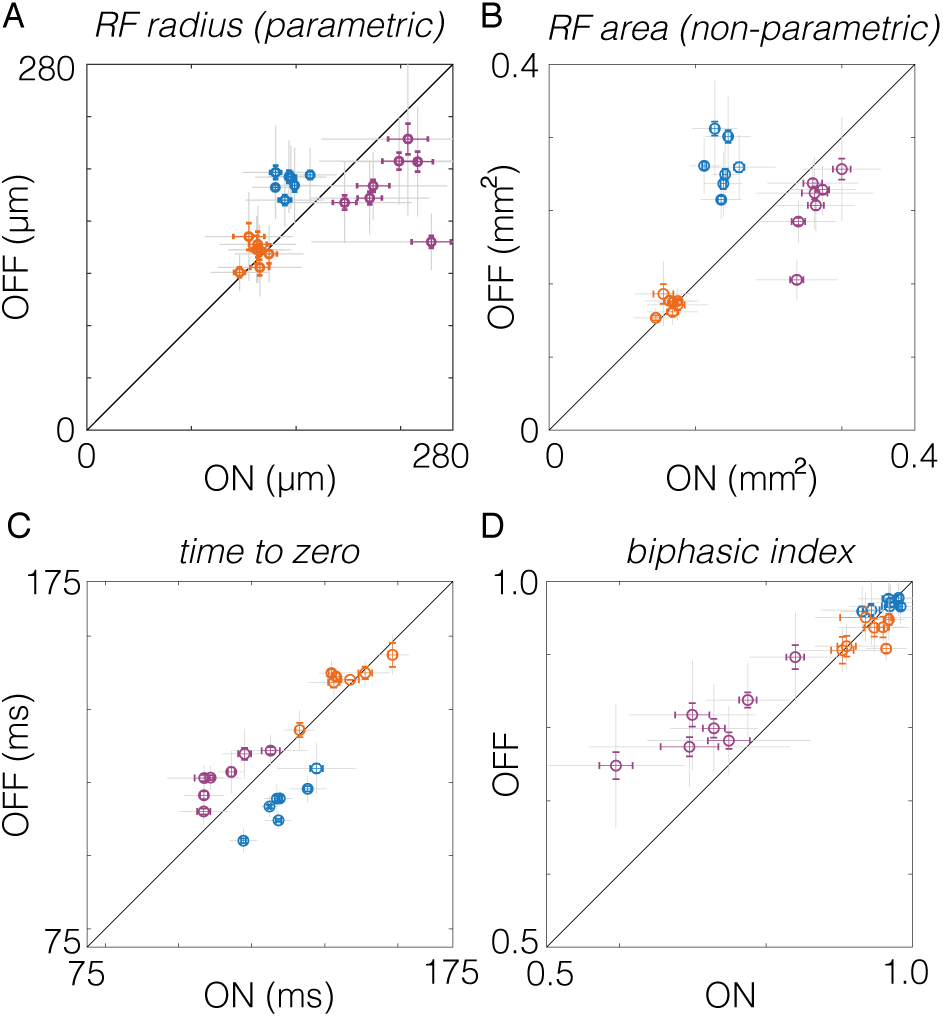
Comparison of spatial and temporal RF properties between ON and OFF RGC pairs. **A.** Spatial RF radii compared between pairs of ON and OFF RGCs. RF radii were derived from a two-dimensional Gaussian fit to the spatial RF. Brisk sustained are purple, brisk transient are blue, and small transient are orange. Each point shows comparison from one retina. Gray error bars show SD, color bars show SE. **B.** Same as A, but compares RF area estimated non-parametrically from the STA (see Materials and Methods). **C.** Comparison of temporal integration estimated from the time to zero of the temporal RF (see Materials and Methods). Comparison of the biphasic index across pairs of ON and OFF RGCs.

#### Parameters of the spatial RF from checkerboard stimuli

The spatial RF diameter (e.g. Figure 6A) was defined as the diameter of a circle with the same area as the 1SD boundary of a two-dimensional Gaussian fit to the RF center (Chichilnisky and Kalmar, 2002; Gauthier et al., 2009). To plot the spatial RF mosaics (e.g. Figure 2A & D), RFs were filtered by convolving with a two-dimensional Gaussian filter with an SD of 0.75 stixels. Contour lines were then linearly interpolated in each RF using a fixed contour equivalent to 1 SD, 0.6065 of the peak (Yu et al., 2017).

## Results

In the following sections, we show the results of a functional classification applied to rat RGCs recorded on a large-scale MEA. This classification yields a natural set of three pairings between ON and OFF RGC types. We analyze the spatiotemporal RF properties and gain among these six cell types and compare the results across ON and OFF pairs.

### The rat retina contains at least three functional pairs of ON and OFF cells

To analyze the RF structure across cell types, we took a serial approach to classifying RGCs (see Materials and Methods). In the first step, direction selective RGCs were separated from other cells based on their responses to gratings drifting in different directions and at different speeds (Figure 1A). In the second step, non-direction-selective RGCs were split into cells with stronger ON or OFF responses (Figure 1B). The dominant response polarity was determined from the spike-triggered average (STA) to a checkerboard stimulus (see Materials and Methods). In the third, fourth and fifth steps, ON and OFF RGCs were serially classified by identifying a small number of response parameters that clustered RGC types. These response parameters included information about the mean firing rates, RF size, and duration/kinetics of temporal integration. This approach yielded three ON and three OFF RGC types.

Across these six RGC types, the classification approach indicated a natural set of three pairs of ON and OFF cell types. For ON and OFF types to be paired, they must resemble one another more than they resemble other cell types, either morphologically (Wassle et al., 1981a) or functionally (Devries and Baylor, 1997). This kind of similarity was indicated by two observations. First, the parameter spaces used to classify ON and OFF RGCs were the same for each pair (Figure 1C-E, steps 3-5). Second, the relative distribution of cells within those parameter spaces were similar for each pair. These two features ensured that the same response properties segregated each pair from all other recorded ON and OFF cells and did so in a similar fashion. These are the core criteria for defining an ON and OFF signaling pair.

Figure 1 The first pair of ON and OFF RGCs (Figure 1C, step 3) were distinguished by their mean spike rate to a drifting grating, the mean response to checkerboard noise, and the ratio between the trough and peak of their temporal RFs. A low trough-to-peak ratio indicates relatively monophasic temporal integration and a ‘sustained’ responses to steps of light. Thus, this first pair of ON and OFF cells exhibited the highest firing rates to drifting gratings and checkerboard noise, relatively sustained responses, and weakly biphasic temporal integration.

After removing this first pair of classified cells, the second pair of ON and OFF RGCs were classified in a new parameter space that compared spatial RF size, duration of temporal integration (time-to-zero), and the mean spike rate to checkerboard noise 1D). For both ON and OFF RGCs, groups of cells exhibited high firing rates to checkerboard noise stimuli, large RFs, and brief temporal integration.

In the final classification step (Figure 1E), the remaining unclassified RGCs were compared in a parameter space consisting of the time-to-zero of the temporal RF, a phasic index calculated on the temporal RF (see Materials and Methods), and the time of the peak in the interspike interval (ISI) distribution. Clusters of ON and OFF cells emerged in these spaces with the briefest ISI peaks, relatively biphasic temporal RFs, and long time-to-zero crossings.

These classification results indicated a set of pairings between ON and OFF RGCs among the cells identified in our MEA measurements. In the subsequent section we examine whether these cells form irreducible types and compare their response properties across a broader range of parameters.

### Each identified ON and OFF cell type forms a mosaic

A hallmark of cell types in the retina is that they tile space morphologically with dendritic fields and functionally with spatial RFs (Wassle and Riemann, 1978; Wassle et al., 1981b; Dacey, 1993; Devries and Baylor, 1997; Novelli et al., 2005; Field and Chichilnisky, 2007). Thus, we tested whether the clusters of ON and OFF cells identified in our serial classification tiled space to form a mosaic-like pattern with their spatial RFs. We measured RGC spatial RFs from STAs to checkerboard noise (see Materials and Methods) (Chichilnisky, 2001; Yu et al., 2017). Plotting the spatial RFs for each RGC type revealed that all six types exhibited a mosaic-like organization (Figures 2A & D). An analysis of the nearest neighbor distributions for RGCs of each type revealed non-random spatial RF organizations for each type across most retinas (Figure 3). Importantly, no information about the spatial location of cells was used at any step of the classification. Thus, the observation of mosaics is a validation that the classification yielded irreducible cell types.

Another feature of RGC types is that response parameters should vary less within a type than across types. Thus, we checked that the temporal RFs (reflecting the temporal integration of visual input) were more similar within a type than across types. Temporal RFs were measured from the STA time courses to checkerboard noise (see Materials and Methods). Plotting the temporal RFs for all six types revealed highly stereotyped temporal integration within a type and distinct temporal integration across types (Figures 2B & E). Finally, we compared (ISI) distributions across types. The ISIs reflect the spiking dynamics of each RGC. Similar to the temporal RFs, the ISI distributions were more similar within a type than across types for both ON and OFF RGCs (Figures 2C & F).

These features of the six RGC types supported the conclusion that each represented an irreducible cell type. Henceforth, we refer to the first pair of classified RGCs (Figure 1C) as ON and OFF brisk sustained RGCs based on their short latency, sustained responses to visual stimuli, and previously used naming conventions (Caldwell and Daw, 1978; Devries and Baylor, 1997; Girman and Lund, 2010; Heine and Passaglia, 2011). Similarly, we refer to the second and third pairs of classified RGCs (Figures 1D-E) as brisk transient and small transient RGCs respectively.

To further test whether the pairings of these types was warranted, we compared the temporal RFs across all six RGC types in a reduced dimensional space defined by principal components analysis (PCA). ON and normalized nearest neighbor distance OFF brisk sustained cells clustered together after accounting for their difference in response polarity (Figures 4A & B). Similarly, ON and OFF brisk transient and ON and OFF small transient cells were more similar to one another, respectively, than to the other identified types. To test that this particular set of pairings was objectively the best three-group association across all six types, we fit a three-Gaussian mixture model to the data, using the first five PCs (Figures 4C & D). The Gaussian mixture model produced an exact match to the three-group description produced by combining ON and OFF cells across brisk sustained, brisk transient, and *PCA on temporal RFs PCA on temporal RFs*< 0.5 small transient cells (compare Figures 4A with C and B with D). This further supports the functional pairings established in the serial classification (Figure 1).

### RGCs with larger spatial integration exhibit briefer temporal integration of visual input

We next compared the spatial and temporal integration of visual input across all six RGC types. Previous studies in primate and cat examining parasol and midget RGCs or alpha and beta RGCs, respectively, have indicated that spatial and temporal integration are inversely related (Frishman et al., 1987; Lee, 1996; Troy and Shou, 2002). Here we examined whether this trend holds in the rodent retina, which has become a dominant model of visual processing (Huberman and Niell, 2011; Sanes and Masland, 2015). Space-time plots of average RFs for each type revealed that types with larger RFs exhibited briefer temporal integration (Figures 5A-F). This relationship held across all seven analyzed retinas (Figure 5G). This comparison assumes that the spatiotemporal integration performed by each RGC is well captured by a single spatial filter and a single temporal filter. We checked the degree of independence between the spatial and temporal RFs: where independence is defined as the STA being well-approximated by the outer product of a spatial and temporal filter (DeAngelis et al., 1993; Golomb et al., 1994; Cai et al., 1997; Cowan et al., 2016). Singular value decomposition revealed that for each of the six RGC types we examined, > 90% of the variance in the STA was captured by the outer product of a single spatial and temporal filter (not shown). These results indicate that for these RGC types, spatiotemporal integration was well approximated by a single spatial and temporal filter. Furthermore, in the rodent retina, as in other species, larger spatial integration implies briefer temporal integration.

### ON-OFF asymmetries in spatial and temporal integration depend on cell type

Previous work has highlighted asymmetries in the size of spatial RFs between ON and OFF cells, with ON cells having larger RFs (Chichilnisky and Kalmar, 2002; Ratliff et al., 2010). To test whether this organization is ubiquitous across ON and OFF pathways in the rodent retina, we compared the size of spatial RFs for each pair of ON and OFF RGC types. Across seven retinas, ON brisk sustained RGCs exhibited larger spatial RFs than OFF brisk sustained RGCs (Figure 6A, purple). However, ON and OFF brisk transient RGCs exhibited the opposite relationship (Figure 6A, blue). Furthermore, ON and OFF small transient cells exhibited nearly identical RF sizes (Figure 6A, orange). These comparisons were based on a two-dimensional Gaussian fit to the spatial RF to identify the radius of a circle with an area equal to that encompassed within one standard deviation of the RF (see Materials and Methods). To test that this result did not depend on a parametric description of the RF, we repeated the comparison for the RF area estimated by the number of stimulus pixels that drove an appreciable change in firing rate for each RGC (see Materials and Methods). Qualitatively, the results were unchanged by the non-parametric analysis (Figure 6B). Thus, previously observed asymmetries do not generalize across cell types.

Previous studies have noted asymmetries in the temporal integration between ON and OFF pathways (Chichilnisky, 2001; Pandarinath et al., 2010) Thus, we next compared the duration of temporal integration between ON and OFF pairs. The duration of the temporal integration was estimated by the time-to-zero between the peak and the trough of the temporal RFs. Consistent with previous results, among brisk sustained RGCs, ON cells exhibited briefer temporal integration than OFF cells (Figure 6C, purple). However, the opposite was observed for brisk transient RGCs (Figure 6C, blue). Similar to the results obtained for spatial RFs, ON and OFF small transient cells exhibited similar durations of temporal integration (Figure 6C, orange).

In addition to the duration of temporal integration, RGCs can differ in the dynamics of integration. A key measure of their temporal dynamics is their biphasic index (a.k.a. degree of transience). For a shift invariant linear system, the biphasic index indicates key properties of temporal filtering (e.g. low-pass vs. bandpass) and it indicates how transient vs. sustained the spiking response will be to a prolonged step in light intensity (Field et al., 2007; Petrusca et al., 2007). Higher biphasic indices indicate more strongly bandpass temporal filtering and more transient light responses. Comparing biphasic indices across ON and OFF pairs, revealed that among brisk sustained RGCs OFF cells exhibited more biphasic temporal integration than ON cells (Figure 6D, purple). However, biphasic indices were similar between ON and OFF cells for brisk and small transient RGCs (Figure 6D, blue and orange). These results indicate that ON-OFF asymmetries in the dynamics of temporal integration are present in some visual pathways, but not all.

### Asymmetries in linearity, gain and SNR among ON and OFF RGC types

The analyses described above compare the spatial and temporal integration of visual input between ON and OFF RGC types. However, these analyses do not reveal differences in spiking output across cell types. The degree of linearity vs. rectification, gain, and signal-to-noise in the spiking output, are all key features dictating the signals provided to downstream brain areas. Previous work has noted that OFF cells are more strongly rectified in their spiking output than ON cells (Chichilnisky and Kalmar, 2002; Zaghloul et al., 2003; Turner and Rieke, 2016), thus pathway asymmetries may extend beyond the integration of sensory input.

To characterize and compare the transformation between visual input and spiking output, we estimated static nonlinearities that relate the filtered visual stimulus to the number of spikes produced by each neuron (Figure 7A) (Chichilnisky, 2001). These static nonlinearities can be thought of as contrast response functions, where contrast is defined as the similarity between the visual stimulus and the spatiotemporal RF.

**Figure 7.**
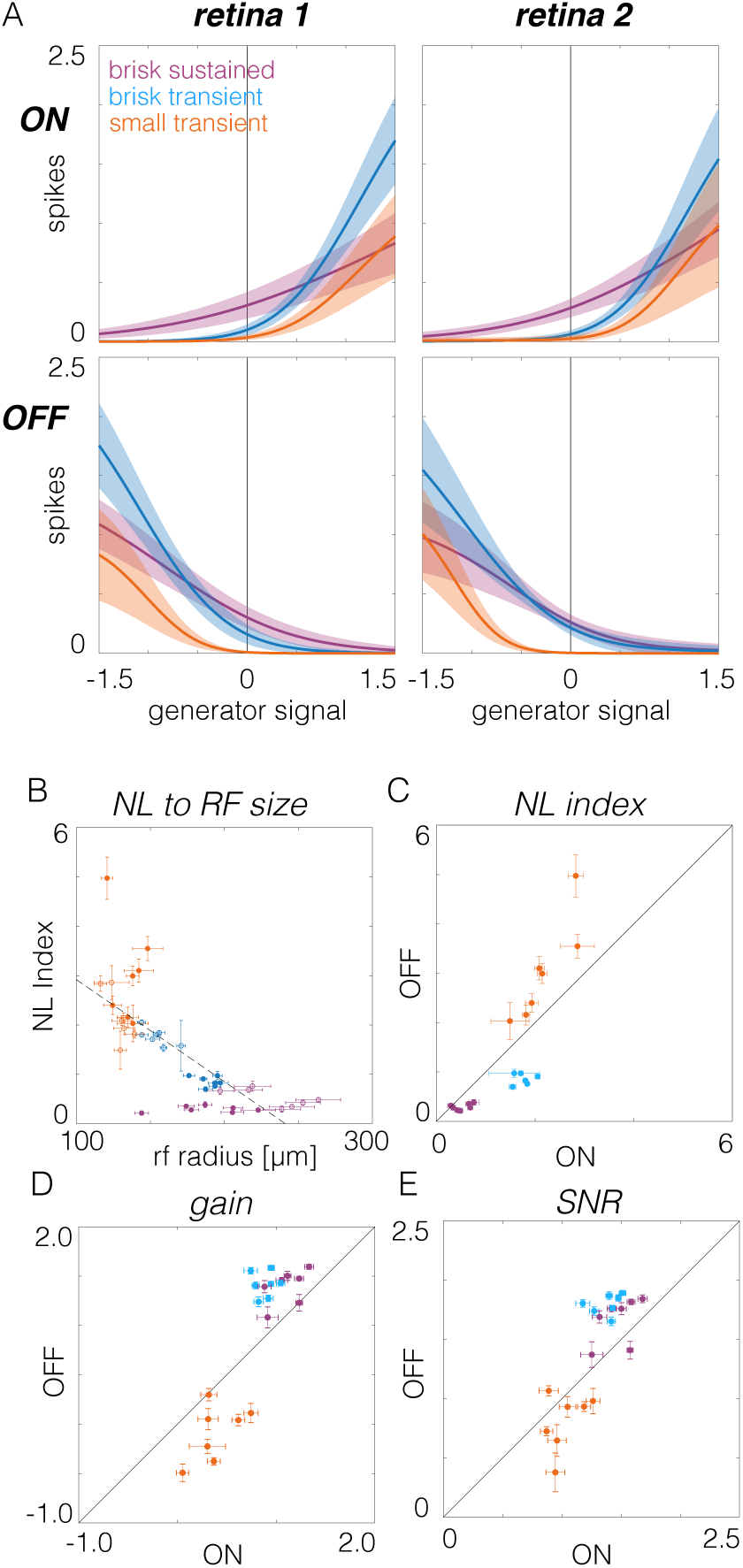
Comparison of contrast response functions across RGC types. **A.** Contrast response functions estimated from the static nonlinearites that relate visual stimuli filtered by the spatiotemporal RF to mean spike counts. Left and right show data from two retinas, top and bottom show ON and OFF RGCs, respectively. **B.** Comparison of nonlinearity index to rf radius. Brisk sustained are purple, brisk transient are blue, and small transient are orange. Filled (open) circles are OFF (ON) cells **C.** Nonlinear index compared between pairs of ON and OFF RGCs. **D.** Gain compared between pairs of ON and OFF RGCs. **E.** Signal-to-noise ratio (SNR) compared between ON and OFF pairs.

ON and OFF brisk sustained RGCs exhibited the most linear contrast response functions (Figure 7A, purple): their spike rates were modulated relatively symmetrically around zero contrast. Brisk transient and small transient cells were progressively more rectified in their spike output (Figure 7A, blue and orange). ON and OFF brisk transient cells exhibited the largest changes in spike rate to large positive or negative contrasts, respectively (Figure 7A, blue).

To relate spiking output to RF properties, we compared RF size to the strength of rectification, as assayed by the NL index, which was computed as the log of the ratios of the slope at the maximum generator signal to the slope at zero generator signal (Chichilnisky and Kalmar, 2002). This comparison revealed that cells with smaller RFs were more rectified in their spiking output than cells with larger RFs (Figure 7B). Because temporal integration was inversely related to spatial integration (Figure 5G), longer temporal integration also implied greater rectification in spike output.

To test for asymmetries in the spiking output of ON and OFF cell types, we first examined NL indices: the logarithm of the ratio of the slope of at the maximum to the slope at zero. For brisk sustained and brisk transient RGCs, ON cells had larger NL indices (greater rectification) than OFF cells (Figure 7C, purple and blue). However, this relationship was reversed for small transient RGCs (Figure 7C, orange). Gain, the log of the slope of the contrast response function at zero contrast, was larger among OFF than ON cells for brisk sustained and brisk transient cells (Figure 7D, purple and blue), but small transient RGCs exhibited the opposite trend (Figure 7D, orange). Finally, the SNR was compared between ON and OFF pathways. The SNR was de-

## Discussion

In this study, we distinguished three functionally matched pairs of ON and OFF cells, which provided an opportunity to test the extent to which ON-OFF asymmetries generalize across a greater range of cell types. This comparison results in an expansion of the diversity of asymmetries present in the mammalian retina. Asymmetries between ON and OFF brisk sustained cells were consistent with previous observations. However, ON and OFF brisk transient cells exhibited asymmetries of opposite polarity and small transient cells exhibited nearly symmetric spatiotemporal integration. Thus, our work alters the conventional view that ON and OFF asymmetries are consistent across diverse RGC types. Below we comment on the method used to classify RGCs in this study, we suggest correspondences to morphologically defined cell types, and we relate the RF organization of RGCs in this study to that observed in other species.

### Functional Classifications of Rodent RGCs

To functionally classify RGCs, we followed the unsupervised classification approach adopted by several previous studies of RGC diversity (Carcieri et al., 2003; Farrow and Masland, 2011; Baden et al., 2016), with the following differences. The first difference was that RGCs were classified using data from individual recordings instead of pooling data across recordings. This reduced the impact of inter-experiment variability which can either blur distinctions between cell types or cause the identification of too many types. Second, relevant response features that distinguished each type were identified before classification. This improved the performance of the Gaussian mixture model because it produced well-separated clusters, thereby minimizing misclassification rates. Only two or three features were selected at each classification step, which kept data requirements for classification relatively low. Third, the classification approach was serial. This mitigated ambiguity in choosing the right number of clusters because each step consisted of fitting just two clusters to the collection of ON cells and two more clusters to the collection of OFF cells (Figures 1C-E). Finally, because many RGCs were recorded in each experiment, this allowed the mosaic arrangement of RFs to provide complementary evidence that the clustered cells were an irreducible type (Wassle et al., 1981b; Devries and Baylor, 1997; Cook and Chalupa, 2000; Field and Chichilnisky, 2007; Anishchenko et al., 2010). Cumulatively, this combination of features facilitated an analysis of the functional organization of six RGC types.

While this approach was reproducible across recordings, it did not classify all recorded cells, nor did it identify all of the functional types. Given an RGC density of ~1500 cells/mm^2^ in the dorsal region of rat retina targeted in these experiments (Danias et al., 2002), 10-15% of RGCs over the electrode array had well-sorted spikes and were tracked across multiple stimulus conditions, requirements for the data analyzed here. Among recorded RGCs, 37+/-3% were not classified because too few cells of other types were sampled. Each stimulus used in this study was presented “full field”, which likely attenuated or silenced spiking in at least some RGC types (e.g. local-edge detectors; (van Wyk et al., 2006; Zhang et al., 2012)). Moreover, only six irreducible RGC types were identified. This falls well short of the ~30 (possibly 40) functionally distinct types that likely exist in the mammalian retina (Field and Chichilnisky, 2007; Völgyi et al., 2009; Sümbül et al., 2014; Sanes and Masland, 2015; Baden et al., 2016). A more complete functional classification of RGC types will be facilitated by using a wider variety of stimuli and developing approaches for recording and spike sorting a higher fraction of RGCs over the MEA (Segev et al., 2004; Prentice et al., 2011; Marre et al., 2012; Yger et al., 2018).

### Correspondences to morphologically defined RGC types

A major goal in retinal research is to generate a complete catalog of RGCs that specifies the correspondences between their function, morphology, and projections to the brain (Sanes and Masland). We did not determine the morphology of the recorded RGCs, however their RF sizes and response kinetics provide some plausible correspondences. The six RGC types examined here all had relatively large RFs and large well-isolated spikes on the MEA. These features indicate large dendritic fields and relatively large somas, suggesting correspondences to the A and C groups of RGCs identified by Sun and colleagues (Sun et al., 2002). The brisk sustained and brisk transient cells likely correspond to the delta and alpha cells identified by Peichl (Peichl, 1989). The ON and OFF small transient cells likely have smaller cell bodies and dendrites in the interior of the IPL because of their transient response properties (Borghuis et al., 2013), suggesting correspondences to the outer and inner B1 RGCs (Huxlin and Goodchild, 1997). We emphasize that these are hypothesized correspondences that require additional experiments to test.

### Diverse contrast response functions across RGC types

The contrast response functions (a.k.a. static nonlinearities) associated with each RGC type differed significantly across the six types we analyzed (Figure 7). Brisk sustained cells were the most linear, while small transient cells were the most rectified in their output. This trend was present across both ON and OFF types. The degree of rectification in RGC output has been largely attributed to rectification in the excitatory synaptic inputs provided by bipolar cells (Zaghloul et al., 2003; Schwartz et al., 2012; Borghuis et al., 2013; Turner and Rieke, 2016). This predicts that the different bipolar cells feeding these distinct RGC types exhibit differing degrees of rectification in their output. These differences are likely shaped by inhibitory amacrine cells (Franke et al., 2017). Importantly, differences in this rectification can play a substantial role in tuning how different cell types respond to natural scenes (Turner and Rieke, 2016).

Several recent studies have also examined the benefit of distinct contrast response functions for encoding, and how these functions can be optimized given constraints imposed by different sources of noise within the retina. One benefit of diverse contrast response functions for encoding is that they could serve to decorrelate a population of neurons responding to complex stimuli. This decorrelation can reduce redundancy in the population code, thereby transmitting the same information with fewer spikes (Barlow, 1961; Vinje and Gallant, 2000; Pitkow and Meister, 2012). Alternatively, different nonlinearities may reflect compensation for noise at different stages of retinal processing to achieve efficient coding (Brinkman et al., 2016). For example, if the dominant source of noise is present before rectification, the most efficient coding is achieved by relatively linear contrast response functions, while more strongly rectified functions are preferred when noise dominates after rectification. Determining how the contrast response functions we observed either serve or constrain the encoding of natural scenes across six parallel processing streams is an important direction for future work.

### Functional asymmetries among ON and OFF pathways

Asymmetries between ON and OFF pathways have been observed across a range of species and contexts. Among primate parasol RGCs, ON cells exhibit larger RFs, briefer temporal integration, and more linear contrast response functions than OFF cells (Chichilnisky and Kalmar, 2002). Some of these asymmetries have been observed in other species and cell types. For example, alpha cells in guinea pigs and brisk sustained cells in rabbits exhibit at least some overlapping asymmetries (Zaghloul et al., 2003; Ratliff et al., 2010; Buldyrev and Taylor, 2013).

The mechanisms that produce some of these asymmetries are clear. For example, systematic differences in spatial RF size likely reflect systematic differences in dendritic field size between some ON and OFF RGC types (Peichl et al., 1987; Dacey and Petersen, 1992; Tauchi et al., 1992; Ratliff et al., 2010). Asymmetries in contrast response functions between ON and OFF alpha cells reflect differences in baseline transmitter release from presynaptic bipolar cells (Zaghloul et al., 2003). Furthermore, differences in intrinsic cellular conductances and synaptic inputs conspire to yield differences in spontaneous firing, spatial nonlinearities, and other properties (Murphy and Rieke, 2006; Margolis and Detwiler, 2007; Zhang and Diamond, 2009; Buldyrev and Taylor, 2013; Turner and Rieke, 2016).

One question raised by these observations is the extent to which these asymmetries meaningfully shape downstream visual processing and perception. Asymmetries in ON and OFF responses originating in the retina clearly influence signals in LGN (Jiang et al., 2015), and shape the responses in primary visual cortex (Yeh et al., 2009; Jin et al., 2011; Komban et al., 2014; Lee et al., 2016). Furthermore, these asymmetries likely underlie psychophysical asymmetries between sensing and processing increments versus decrements of light (Pons et al., 2017).

Several studies have indicated that ON-OFF asymmetries are optimizations to the statistics of natural scenes. First, a theoretical analysis indicates that the division of processing ON and OFF signals transmits more information with fewer spikes than alternative encoding strategies (Gjorgjieva et al., 2014). Second, the observation that at least some OFF pathways have smaller RFs than ON cells may allow the retina to transmit more information about natural scenes, which exhibit more regions of relative darkness (Ratliff et al., 2010). Similarly, several asymmetries can be predicted by applying efficient coding theory to natural scenes (Karklin and Simoncelli, 2011; Doi et al., 2012).

Given previous work suggests that natural scenes and efficient coding can predict one set of asymmetries (e.g. ON cells having larger spatial RFs than OFF), why do different pathways exhibit different asymmetries? One possibility comes from a recent analysis of the spatial frequency distribution of light and dark asymmetries in natural scenes (Cooper and Norcia, 2015). This work shows that intensity distributions are skewed toward darker values at low spatial frequencies, but not at higher spatial frequencies. This may explain why cell types with the smallest RFs in this study exhibited nearly equivalent spatiotemporal integration (Figures 6A & B). Two other considerations may be important as well. First, previous analyses of natural scenes have largely focused on static images, not on natural movies, or movies that consider head and eye movements. These temporal dynamics may interact with the differences in temporal integration across RGC types to cause different asymmetries to be optimal. Second, previous analyses have largely focused on just two pathways, one ON and one OFF (Karklin and Simoncelli, 2011). It is unclear that the conclusions for encoding natural scenes under this context will generalize if a system has more pathways to utilize for encoding visual scenes. To resolve these possibilities, a more complete analysis of the interactions between the natural movies (including head and eye movements; (Wallace et al., 2013)) and the spatiotemporal dynamics of RGC RFs will be required.

## Acknowledgements

We thank Jon Cafaro, Lindsey Glickfeld, Fred Rieke, and Kiersten Ruda for comments on the manuscript, and Teleza Westbrook, Alexander Sher, and Alan M. Litke for technical support. National Institutes of Health and National Eye Institutes R01s EY024567 (G.D.F) EY021271 (EJC), EY017992 (EJC), P30 EY019005 (EJC), the Karl Kirchgessner Foundation (G.D.F), the Whitehall Foundation (G.D.F), and the Whitehead Scholars Program (G.D.F).

## Author Contributions

Conceptualization, S.R., E.J.C., G.D.F.; Data Collection, D.A, M.G., S.R., G.D.F.; Data Analysis., S.R., D.A., G.D.F., Writing & Editing, S.R., E.J.C., G.D.F.; Supervision, G.D.F.; Funding Acquisition, E.J.C., G.D.F.

